# Assessing introgressive hybridization in roan antelope (*Hippotragus equinus*): Lessons from South Africa

**DOI:** 10.1101/569830

**Authors:** Anna M van Wyk, Desiré L Dalton, Antoinette Kotzé, J. Paul Grobler, Prudent S. Mokgokong, Anna S Kropff, Bettine Jansen van Vuuren

## Abstract

Biological diversity is being lost at unprecedented rates, with admixture and introgression presenting major threats to species’ conservation. To this end, our ability to accurately identify introgression is critical to manage species, obtain insights into evolutionary processes, and ultimately contribute to the Aichi Targets developed under the Convention on Biological Diversity. A case in hand concerns roan antelope, one of Africa’s most iconic large mammal species. Despite their large size, these antelope are sensitive to habitat disturbance and interspecific competition, leading to the species being listed as Least Concern but with decreasing population trends, and as extinct over parts of its range. Molecular research identified the presence of two evolutionary significant units across their sub-Saharan range, corresponding to a West African lineage and a second larger group which includes animals from East, Central and Southern Africa. Within South Africa, one of the remaining bastions with increasing population sizes, there are a number of West African roan antelope populations on private farms, and concerns are that these animals hybridize with roan that naturally occur in the southern African region. We used a suite of 27 microsatellite markers to conduct admixture analysis. Our results unequivocally indicate evidence of hybridization, with our developed tests able to accurately identify F1, F2 and non-admixed individuals at threshold values of *qi* = 0.20 and *qi* = 0.15, although further backcrosses were not always detectable. Our study is the first to confirm ongoing hybridization in this iconic African antelope, and we provide recommendations for the future conservation and management of this species.

## Introduction

The increased rate of human-driven global change is a major threat to biodiversity [1]. Factors such as climate change, habitat fragmentation, and environmental degradation are influencing the distribution and abundance of species, often in ways that are impossible to predict [2]. Thus, a central theme in conservation biology is how best to manage for species persistence under rapidly changing and often unpredictable conditions. When faced with environmental change, species may persist by moving (or being moved) to track suitable environments. Although there is sufficient evidence to suggest that species notably alter their ranges [3], facilitation of such movement for larger vertebrate species (through the creation of habitat corridors, transfrontier parks or translocations) often place insurmountable burdens on conservation agencies that are ultimately responsible for the management of these populations. Notwithstanding, signatory countries to the Convention on Biological Diversity have an obligation to manage and protect biodiversity, as also set out more recently in the Aichi Biodiversity Targets.

Admixture and introgression are major threats to species conservation (these threats are dealt with specifically under Aichi Target 13; see https://www.cbd.int/sp/targets/). The ability to accurately identify introgression is critical to the management of species [4–9], and may provide unprecedented insights into evolutionary processes. Although admixture, or even genetic rescue, may have beneficial outcomes through the introduction of new alleles into small or isolated populations, it can lead to outbreeding depression essentially disrupting locally adapted gene-complexes [10–13]. Because of the movement of animals (either natural or human-facilitated), admixture and the effects thereof become increasingly more important to understand and manage.

Roan antelope (*Hippotragus equinus*) is one of Africa’s most iconic large antelope species. It has a sub-Saharan range, is a water-dependant species, and prefers savanna woodlands and grasslands. [14] recognised six subspecies namely *H. e. equinus, H. e. cottoni, H. e. langheldi, H. e. bakeri, H. e. charicus*, and *H. e. koba* based on morphological analyses. However, subsequent genetic studies by [15] and [16] provided less support for these subspecies designation. Although the [15] study included relatively few specimens (only 13 animals were available at the time), [16] analyzed 137 animals sampled from across the range (the only subspecies not included in this study was *H. e. bakeri*) for both the mtDNA control region and eight microsatellite markers. Both the mtDNA control region and microsatellite data provided strong support for a separation between the West Africa population (corresponding to the *H. e. koba* subspecies) and those from East, Central and Southern Africa (representing the *H. e. equinus, H. e. langheldi*, and *H. e. cottoni* subspecies). Although some differentiation between East, Central and Southern African roan antelope was evident from the mtDNA data, the different subspecies did not form monophyletic groups, with no differentiation observed for the microsatellite data. The placement of the two specimens from Cameroon (corresponding to the *H. e. charicus* subspecies) were unclear, and the small sample size precluded robust analyses. Based on these results, [16] argued that two evolutionary significant units should be recognized for roan antelope in Africa, corresponding to a West African lineage and an East, Central and Southern African lineage.

Roan antelope is listed as Least Concern, but with decreasing population sizes, notably in East and Southern Africa [17]. In Southern Africa, roan antelope numbers have dramatically declined in Botswana, Namibia and Zimbabwe and these animals have been eliminated from large parts of their former range including Angola and Mozambique [18]. Within South Africa, roan antelope numbers in reserves and protected areas are critically low, with the majority of animals residing under private ownership on game farms. Indeed, the estimated population size of wild and naturally occurring roan antelope in protected areas in South Africa is less than 300 animals [19], yet indications are that roan antelope is thriving on private land. Current estimates suggest that at least 3,500 individuals are managed on private farms [20], with numbers increasing due to these animals being considered an economically important species by the South African wildlife industry. In the 1990s, a number of roan antelope (approximately 40) was imported into South Africa under permit from West Africa. Subsequent to their import, and based on DNA evidence [16], an embargo was placed on the trade of West African animals in South Africa. Recent anecdotal evidence suggested that animals of West African decent was being traded in (based on mitochondrial haplotypes; Jansen van Vuuren, pers. comm.), thereby presenting a real and significant threat to the genetic integrity of roan antelope in South Africa, notwithstanding legislation prohibiting it. Furthermore, animals are sometimes being exported to other Southern African countries, further endangering regional gene pools.

Our aim here is to expand on the limited and non-specific suite of microsatellite markers employed by [16] to specifically test the validity of these anecdotal reports of trade in West African roan. Also, we assessed the ability of these markers to discriminate between non-admixed animals and hybrid offspring (F2, F3, and F4). Our results will not only confirm whether suggestions of hybridization are true, but will also provide a valuable tool to ensure genetic integrity in the conservation of roan antelope in Southern Africa.

## Materials and Methods

### Sampling

Blood, tissue or hair material was obtained from private breeders and game farm owners throughout South Africa (Table 1). Reference samples were selected from the [16] study and represent animals of confirmed provenance. A total of 32 West African roan antelope (populations from three farms in Limpopo Province, South Africa), and 98 animals representing the East, Central and Southern African ESU (populations from two farms in the Northern Cape and North West provinces, South Africa) were included. In addition, eight known hybrids and 15 putative hybrids were included in this study (Table 1), provided to us by game owners that legally had West African roan on their farms. Ethical approval was obtained from the Animal Research Ethics Committee, University of the Free State, South Africa (UFS-AED2017/0010) and the NZG Research Ethics and Scientific Committee (NZG/RES/P/17/18). Samples were stored in the NZG Biobank and access for research use of the samples was approved under a Section 20 permit from the Department of Agriculture, Forestry and Fisheries, South Africa (S20BB1917).

**Table 1.**
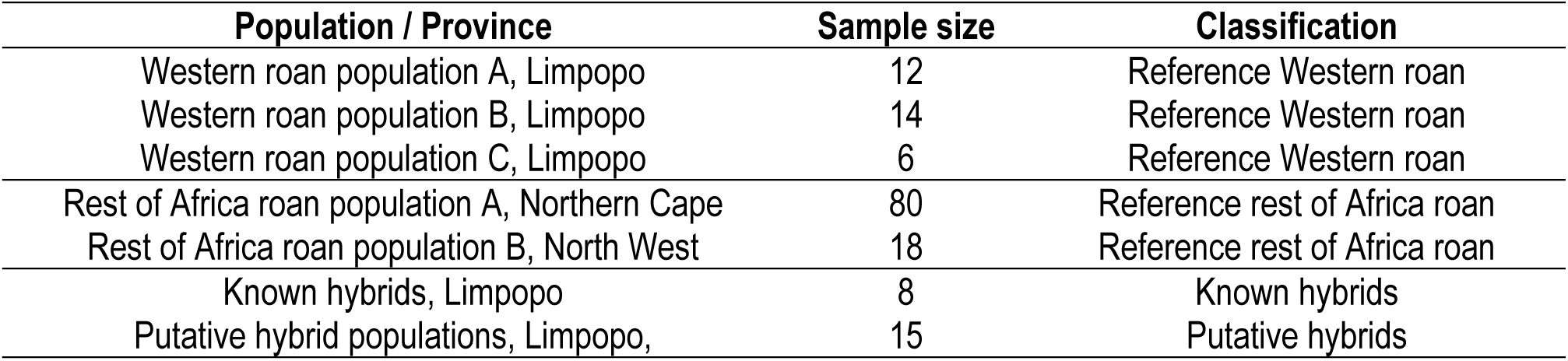
List of roan antelope (*Hippotragus equinus*) samples.

| Population / Province | Sample size | Classification |
| --- | --- | --- |
| Western roan population A, Limpopo | 12 | Reference Western roan |
| Western roan population B, Limpopo | 14 | Reference Western roan |
| Western roan population C, Limpopo | 6 | Reference Western roan |
| Rest of Africa roan population A, Northern Cape | 80 | Reference rest of Africa roan |
| Rest of Africa roan population B, North West | 18 | Reference rest of Africa roan |
| Known hybrids, Limpopo | 8 | Known hybrids |
| Putative hybrid populations, Limpopo, | 15 | Putative hybrids |

### Microsatellite markers

We selected nine cross-species microsatellite markers (HN60, HN02, HN17, HN27, HN113, HN58, HN09, HN12 and HN13) that were previously characterised in sable antelope (*Hippotragus niger*) by Vaz Pinto [7] and 12 cross-species microsatellite markers (BM3517, BM203, SPS113, BM1818, OARFCB304, CSSM19, ILST87, BM719, BM757, OARCP26, OARFCB48, INRA006) that were developed for domestic livestock [21–28]. In addition, species-specific microsatellite markers were developed from non-admixed East, Central and Southern African roan using a Next Generation Sequencing approach. The Nextera^®^ DNA Sample Preparation Kit (Illumina, Inc., San Diego, California, USA) was used to create a paired-end library followed by sequencing on the MiSeq™ sequencer (Illumina, Inc., San Diego, California, USA) using 2 x 300 bp chemistry. Library construction and sequencing was carried out at the Agricultural Research Council Biotechnology Platform (Onderstepoort, Gauteng, South Africa). FastQC version 0.11.4 [29] and Trimmomatic version 0.36 [30] were used for quality control of the raw sequence reads. Tandem Repeat Finder version 4.09 [31] was used to search the remaining reads for microsatellite motifs and Batchprimer3 software [32] was used to design primer pairs flanking the repeat regions.

### Polymerase Chain Reaction (PCR) and genotyping

DNA extractions were performed using the Qiagen DNeasy^®^ Blood and Tissue Kit (Qiagen GmbH, Hilden, Germany) following the manufacturer’s protocols. Polymerase Chain Reaction (PCR) amplification was conducted in 12.5 μl reaction volumes consisting of AmpliTaq^®^ DNA polymerase (Roche Molecular Systems, Inc) forward and reverse primers (0.5 μM each), and 50 ng genomic DNA template. The conditions for PCR amplification were as follows: 5 min at 95°C denaturation, 35 cycles for 30 sec at 95°C, 30 sec at 50-62°C (primer-specific annealing temperatures) and 30 sec at 72°C, followed by extension at 72°C for 10 min in a T100™ Thermal Cycler (Bio-Rad Laboratories, Inc. Hercules, CA, USA). PCR products were run against a Genescan™ 500 LIZ™ internal size standard on an ABI 3130 Genetic Analyzer (Applied Biosystems, Inc., Foster City, CA, USA). Samples were genotyped using GeneMapper v. 4.0 software (Applied Biosystems, Inc., Foster City, CA, USA).

### Genetic diversity

Understanding the diversity within groups provide valuable information to identify hybrid individuals. To this end, genetic diversity was evaluated for each group separately (the two different ESUs, known hybrids, and putative hybrids). MICRO-CHECKER [33] was used to detect possible genotyping errors, allele dropout and null alleles. The mean number of alleles per locus (A), allelic richness (AR), observed heterozygosity (Ho), unbiased heterozygosity (Hz = expected heterozygosity adjusted for unequal sample sizes) [34] and number of private alleles per reference group (N_P_) was calculated with GenAlEx 6.5 [35,36]. Arlequin 3.5 [37,38] was used to test for deviations from expected Hardy-Weinberg (HW) proportions of genotypes (Markov Chain length of 105 and 100,000 dememorization steps) and to evaluate loci for gametic disequilibrium (with 100 initial conditions followed by ten permutations, based on the exact test described by Guo and Thompson [39]. Associated probability values were corrected for multiple comparisons using Bonferroni adjustment for a significance level of 0.05 [40]. In addition, to determine the discriminatory power of the combined loci, the P_ID_ was calculated using GenAlEx [35,36]. Finally, inbreeding (F_IS_) and average pairwise relatedness between individuals within populations was calculated using the R package Demerelate version 0.9-3 (using 1,000 bootstrap replications) [41].

### Population structure and admixture analysis

To estimate the degree of genetic differentiation between populations, we performed an analysis of molecular variance (AMOVA) and conducted pairwise F_ST_ comparisons among populations in ARLEQUIN version 3.5 [37,38]. We used two approaches to assess population structure, namely a Bayesian clustering approach implemented in STRUCTURE version 2.3.4 [42–44] and a Principal Component Analysis (PCA). STRUCTURE was used for the identification of genetic clusters and individual assignment of non-admixed animals as well as putative hybrid individuals and was run using a model that assumes admixture, correlated allele frequencies and without prior population information for five replicates each with K = 1 – 6, with a run-length of 700,000 Markov Chain Monte Carlo repetitions, following a burn-in period of 200,000 iterations. The five values for the estimated ln(Pr (X|K)) were averaged, from which the posterior probabilities were calculated. The K with the greatest increase in posterior probability (ΔK) [45] was identified as the optimum number of sub-populations using STRUCTURE HARVESTER [46]. The membership coefficient matrices (Q-matrices) of replicate runs for the optimum number of sub-populations was combined using CLUMPP version 1.1.2 [47] with the FullSearch algorithm and G′ pairwise matrix similarity statistics. The results were visualized using DISTRUCT version 1.1 [48]. From the selected K value, we assessed the average proportion of membership (*qi*) of the sampled populations to the inferred clusters. Individuals (parental or admixed classes) were assigned to the inferred clusters using an initial threshold of *qi* > 0.9 [49]. PCA for the complete data set was achieved using the R package Adegent version 2.1.1 [50].

### Maximizing the accuracy of assignments

To determine which threshold Q-value (hybridization or admixture index from clustering algorithms like STRUCTURE) would maximize the accuracy of assignment, simulated genotypes were created using HYBRIDLAB [51]. Genotypes of non-admixed Western roan antelope, and animals from East, Central and Southern Africa (n =30) with *qi* > 0.90 (from STRUCTURE-based analysis) were used a parental (P1) populations to create the simulated hybrid genotypes (see [9]). A dataset consisting of 180 individuals were created consisting of 30 each belonging to non-admixed Western roan antelope, non-admixed Eastern, Central and Southern roan antelope, F1 hybrids, F2 hybrids, backcrosses of F1 with Western roan (BC-Western roan) and backcrosses of F1 with rest of Africa roan antelope (BC-rest of Africa roan). The simulated dataset was analysed with STRUCTURE version 2.3.4 [42–44] using the admixed model, correlated allele frequencies and without prior population information for five replicates each with K = 1 – 2, a run-length of 700,000 Markov Chain Monte Carlo repetitions and a burn-in period of 200,000 iterations.

## Results

### Species-specific microsatellite markers

In this study, species specific microsatellite markers were successfully developed using DNA extracted from non-admixed roan antelope (i.e., animals of known provenance). Read lengths of 2 x 301 bp (2 x 3,306,938) were obtained and after trimming, the remaining reads ranged from 180 to 200 bp (2 x 1,596,026). A total of 14 unique loci were identified, of these only six were polymorphic and consistently amplified animals from both ESUs (Table 2).

**Table 2.**
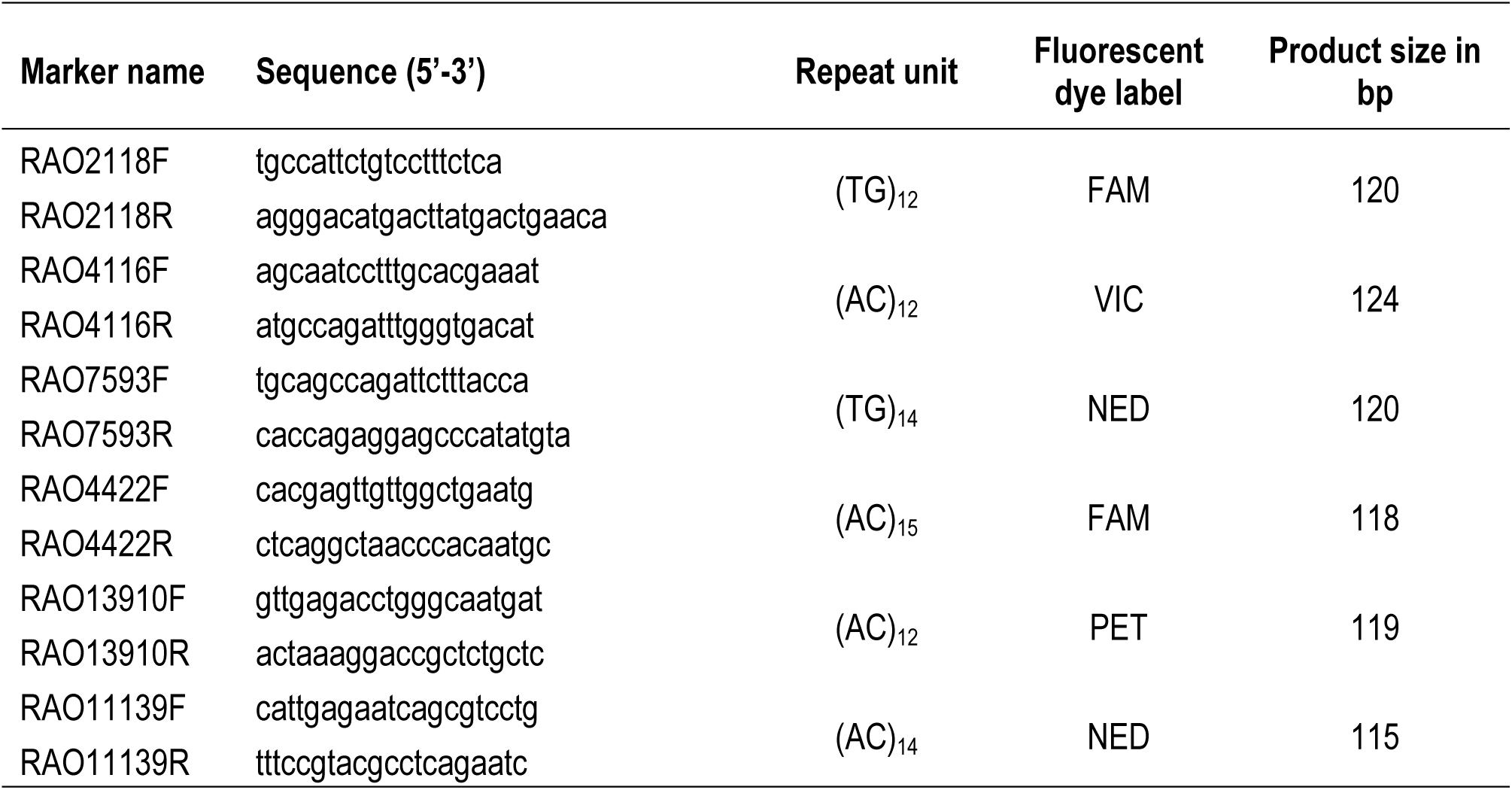
List of six species-specific microsatellite loci developed in *Hippotragus equinus*: F = forward primer; R = reverse primer; bp = base pairs. GenBank accession numbers are MN699986-MH699992.

### Genetic differentiation and admixture analysis

The final dataset included 27 microsatellite loci that yielded a total of 267 alleles, with the number of alleles ranging from 3 to 17 per locus. A total of 27 alleles were unique to the West African roan group, while 27 were found exclusively in the East, Central and Southern African group (Table 3). An analysis of molecular variance (AMOVA) unequivocally retrieved the two distinct groups (corresponding to the two ESUs reported by Alpers [16]; F_ST_ = 0.165, P < 0.001), validating our two reference groups. Principle component analysis similarly revealed a clear separation between the West African versus East, Central and Southern Africa roan (Fig 1A). The two distinct genetic clusters (K = 2) was supported by the Bayesian assignment analysis (Fig 1B, S1 Fig). West African versus East, Central and Southern African roan antelope were assigned to two distinct clusters with individual coefficient of membership (*qi*) for non-admixed Western roan *qi* > 0.881 and for non-admixed East, Central and Southern Africa roan *qi* > 0.883. With regards to known hybrids, six of the eight known hybrids were confirmed as hybrids, with two hybrids being identified as non-admixed Western roan (*qi* = 0.9664 and *qi* = 0.9510, respectively). Analysis of putative hybrids identified four out of 15 animals as hybrid (27%).

**Table 3.** Private alleles in loci and allele frequency in Western and rest of Africa roan.

| Population | Locus | Allele | Frequency | Population | Locus | Allele | Frequency |
| --- | --- | --- | --- | --- | --- | --- | --- |
| Western roan | BM203 | 230 | 0.040 | Rest of Africa roan | BM203 | 240 | 0.005 |
| Western roan | Oarc26 | 146 | 0.031 | Rest of Africa roan | BM719 | 169 | 0.005 |
| Western roan | Oarc26 | 148 | 0.047 | Rest of Africa roan | BM719 | 177 | 0.074 |
| Western roan | OARFCB48 | 176 | 0.078 | Rest of Africa roan | Oarc26 | 118 | 0.010 |
| Western roan | BM1818 | 280 | 0.089 | Rest of Africa roan | Oarc26 | 124 | 0.015 |
| Western roan | BM757 | 180 | 0.031 | Rest of Africa roan | BM1818 | 256 | 0.058 |
| Western roan | ILST87 | 153 | 0.018 | Rest of Africa roan | BM1818 | 278 | 0.016 |
| Western roan | ILST87 | 159 | 0.018 | Rest of Africa roan | BM1818 | 282 | 0.068 |
| Western roan | RAO4422 | 115 | 0.017 | Rest of Africa roan | BM1818 | 288 | 0.037 |
| Western roan | RAO4422 | 129 | 0.017 | Rest of Africa roan | INRA006 | 117 | 0.077 |
| Western roan | RAO4422 | 137 | 0.050 | Rest of Africa roan | INRA006 | 123 | 0.056 |
| Western roan | RAO4422 | 141 | 0.050 | Rest of Africa roan | INRA006 | 125 | 0.077 |
| Western roan | RAO4422 | 149 | 0.017 | Rest of Africa roan | INRA006 | 127 | 0.337 |
| Western roan | RAO4422 | 151 | 0.033 | Rest of Africa roan | OARFCB304 | 115 | 0.016 |
| Western roan | RAO4422 | 155 | 0.033 | Rest of Africa roan | OARFCB304 | 127 | 0.005 |
| Western roan | RAO4422 | 159 | 0.017 | Rest of Africa roan | ILST87 | 121 | 0.005 |
| Western roan | RAO13910 | 115 | 0.031 | Rest of Africa roan | ILST87 | 127 | 0.005 |
| Western roan | RAO4116 | 126 | 0.047 | Rest of Africa roan | RAO13910 | 141 | 0.040 |
| Western roan | HN02 | 186 | 0.063 | Rest of Africa roan | RAO11139 | 102 | 0.010 |
| Western roan | HN17 | 202 | 0.109 | Rest of Africa roan | RAO11139 | 104 | 0.026 |
| Western roan | HN58 | 124 | 0.031 | Rest of Africa roan | RAO11139 | 108 | 0.072 |
| Western roan | HN58 | 144 | 0.016 | Rest of Africa roan | RAO4116 | 112 | 0.086 |
| Western roan | HN09 | 152 | 0.047 | Rest of Africa roan | HN09 | 168 | 0.005 |
| Western roan | HN09 | 180 | 0.031 | Rest of Africa roan | HN09 | 173 | 0.005 |
| Western roan | HN09 | 194 | 0.031 | Rest of Africa roan | HN12 | 185 | 0.005 |
| Western roan | HN12 | 171 | 0.032 | Rest of Africa roan | HN12 | 195 | 0.005 |
| Western roan | HN12 | 193 | 0.032 | Rest of Africa roan | HN13 | 184 | 0.025 |

**Fig 1.**
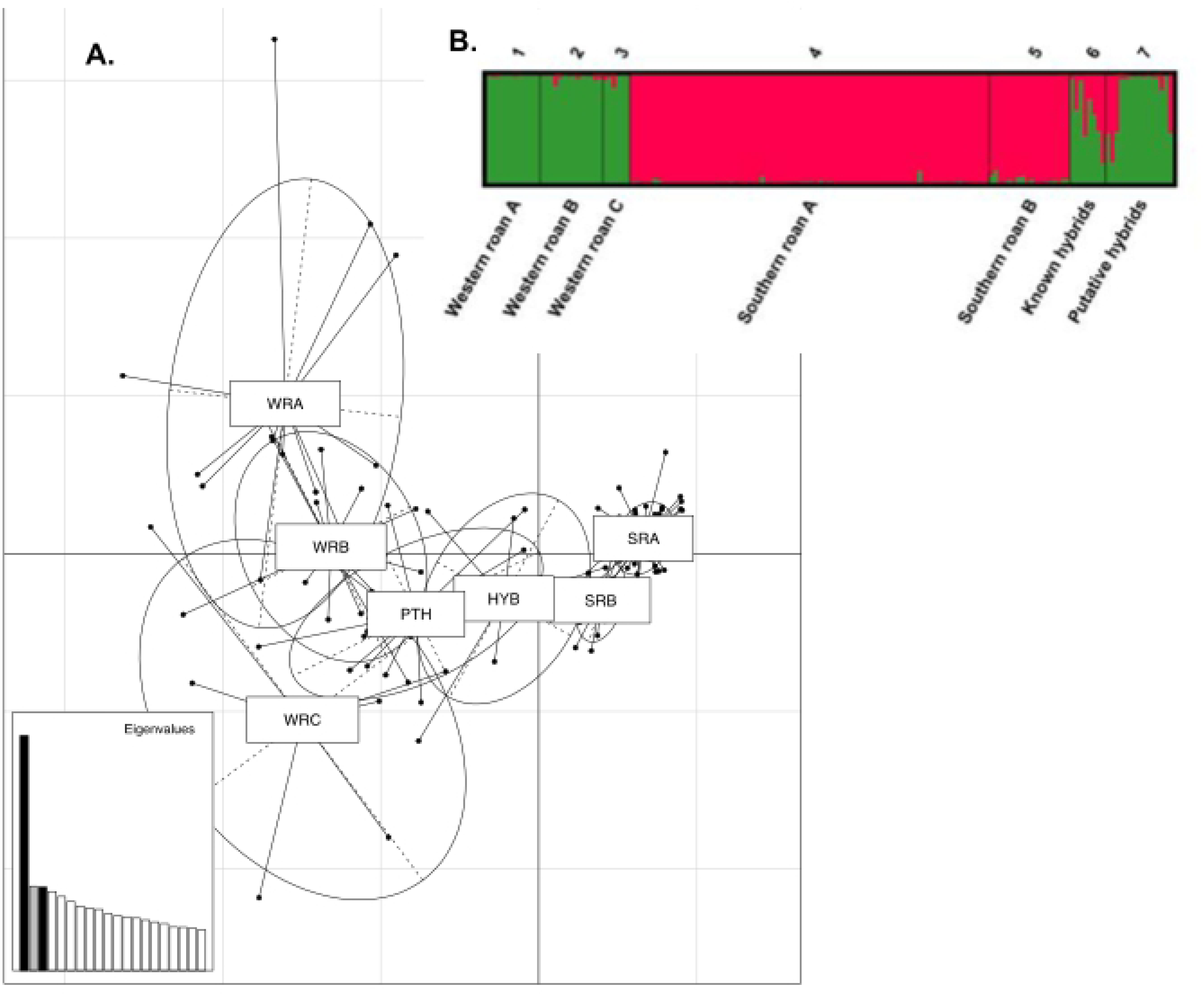
Genetic differentiation analysis between populations based on (A) Principal Component Analysis (PCA) and (B) STRUCTURE analysis (performed with K = 2) of Western roan, rest of Africa roan, known hybrids and putative hybrids. WRA = Western roan A, WRB = Western roan B, WRC = Western roan C, SRA = rest of Africa A, rest of Africa B, HYB = known hybrids, PTH = putative hybrids.

On South African farms, game owners often employ selective breeding to achieve specific outcomes. For example, hybrid animals may be backcrossed with pure roan to selectively breed hybrid lineages back to pure; in theory this can be achieved in N = 4 generations. We wanted to assess whether our markers are able to detect backcrossed animals, especially in the F3 and F4 generations. In this study, we created a simulated dataset to maximize the accuracy of assignment to distinguish between the two non-admixed groups (West Africa versus East, Central and Southern roan antelope), F1 hybrids, F2 hybrids, F1 BC-Western roan and F1 BC-rest of Africa roan. STRUCTURE analysis of simulated genotypes generated by HYBRIDLAB indicated that all (100%) of the West African roan versus East, Central and Southern Africa roan, F1 and F2 genotypes were correctly assigned at thresholds of *qi* > 0.80 and *qi* > 0.85 (Table 4). At a threshold value of *qi* > 0.90, all F1, F2 hybrid and the East, Central and Southern Africa roan were correctly assigned, however, 20% of the non-admixed Western roan would be incorrectly identified as hybrid origin. At a threshold value of *qi* > 0.95, all F1 and F2 hybrid individuals would be correctly assigned, however, 40% of non-admixed Western roan and 7% of the East, Central and Southern African roan would be incorrectly identified as hybrid. Our ability to distinguish non-admixed roan from backcrossed individuals may be problematic in some instances with correct assignment of backcrossed Western roan individuals varying from 40% at *qi* > 0.80 to 97% at *qi* > 0.95, and backcrossed East, Central and Southern African roan individuals varying from 53% at *qi* > 0.80 to 97% at *qi* > 0.95. Based on the simulation results, the threshold *q*-value of *qi* >0.85 was selected for analysis of the non-admixed parental populations, known hybrids and putative hybrids.

**Table 4:**
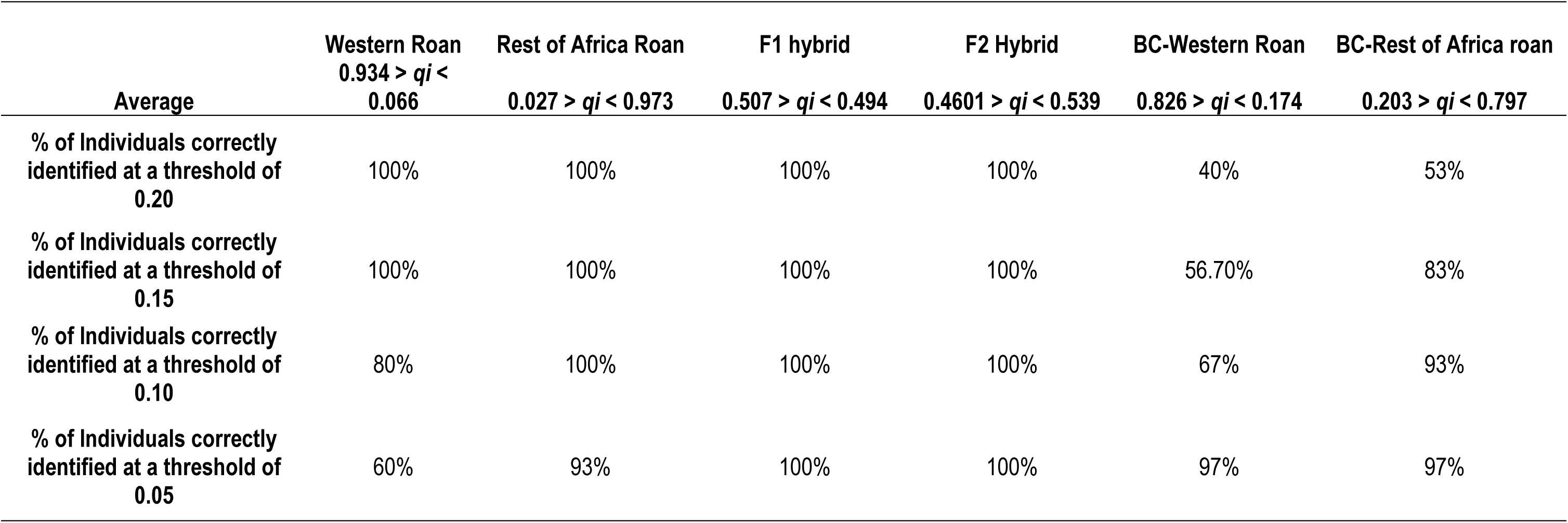
Percentage of individuals correctly identified at different threshold values.

| Average | Western Roan<br>0.934 > *qi* <<br>0.066 | Rest of Africa Roan<br>0.027 > *qi* < 0.973 | F1 hybrid<br>0.507 > *qi* < 0.494 | F2 Hybrid<br>0.4601 > *qi* < 0.539 | BC-Western Roan<br>0.826 > *qi* < 0.174 | BC-Rest of Africa roan<br>0.203 > *qi* < 0.797 |
| --- | --- | --- | --- | --- | --- | --- |
| % of Individuals correctly identified at a threshold of 0.20 | 100% | 100% | 100% | 100% | 40% | 53% |
| % of Individuals correctly identified at a threshold of 0.15 | 100% | 100% | 100% | 100% | 56.70% | 83% |
| % of Individuals correctly identified at a threshold of 0.10 | 80% | 100% | 100% | 100% | 67% | 93% |
| % of Individuals correctly identified at a threshold of 0.05 | 60% | 93% | 100% | 100% | 97% | 97% |

### Genetic diversity and relatedness

Deviations from HWE equilibrium were not consistent across populations, with significant deviations from HWE being observed only in the East, Central and Southern African roan populations. In the East, Central and Southern Africa roan population A (Northern Cape Province), 11 loci (BM3517, BM719, OARFCB48, CSSM19, BM1818, BM757, SPS113, INRA006, OARFCB304, RAO4116 and HN27) deviated from HWE. In addition, two loci (BM3517 and SPS113) deviated from HWE in East, Central and Southern Africa roan population B (North West Province) following Bonferroni correction. These markers indicated significant heterozygote deficit in the respective populations with H_o_ values lower than H_e_ values, which may be an indication of the presence of possible null alleles. However, null alleles were only observed in six markers (BM3517, BM719, SPS113, INRA006, RAO4116 and HN27) from the East, Central and Southern African roan group. Significant linkage disequilibrium (LD) was also observed only in the East, Central and Southern African group. These departures from equilibrium may be because of substructure in this group (see [16], which described three mitochondrial DNA groups within this larger ESU), or because of inbreeding. To further investigate the possible causes of heterozygote deficiency, we estimated the overall inbreeding coefficient per population with positive estimates only being observed in the East, Central and Southern African roan group (F = 0.102). In addition, analysis of the overall population relatedness was conducted, as mating among close relatives may cause heterozygote deficiency. As shown in Fig 2, the overall population relatedness was higher in the East, Central and Southern African roan group (average = 74%) compared to the West African animals (average = 39%).

**Fig 2.**
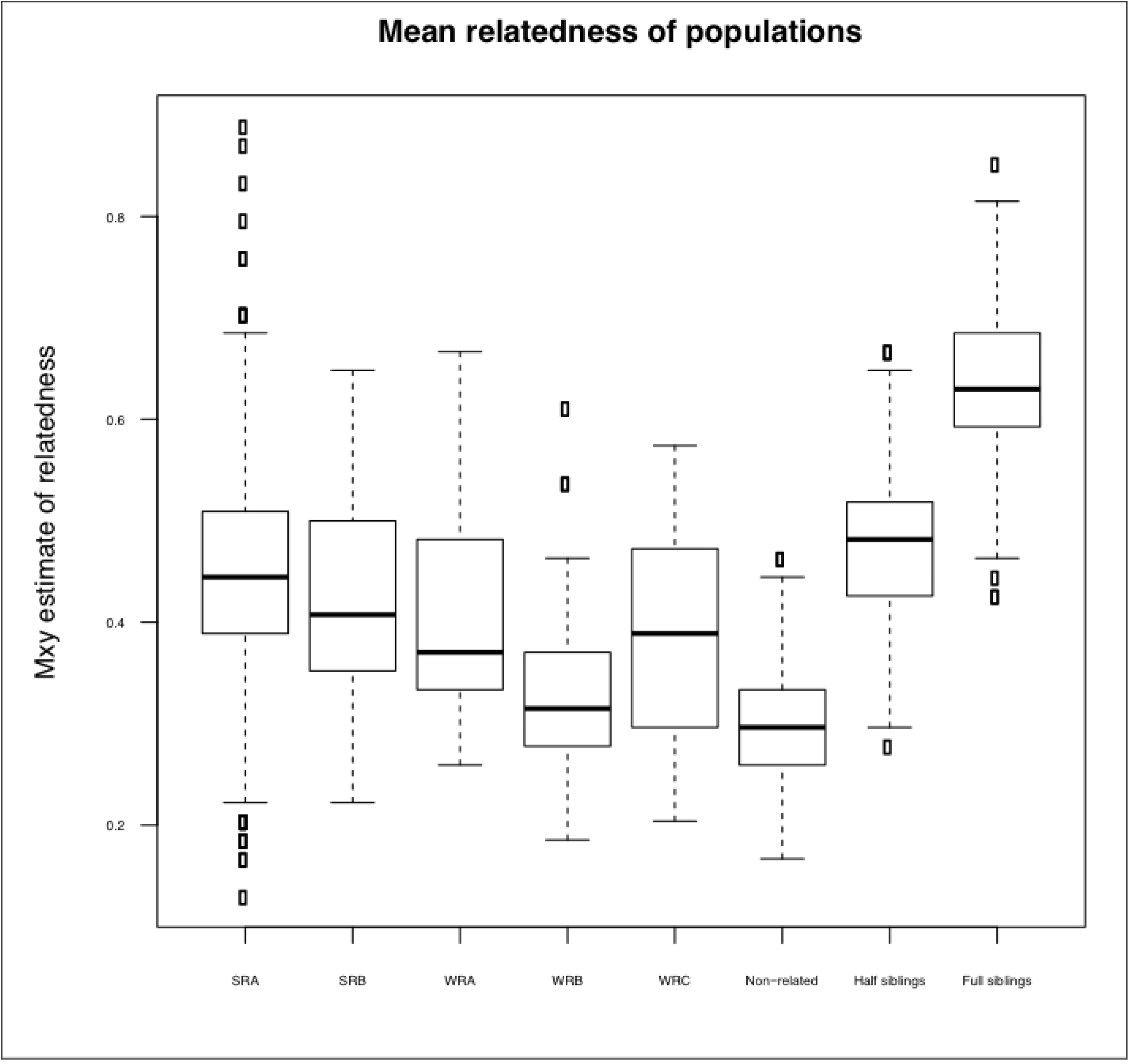
Mean relatedness of rest of Africa roan and Western roan. WRA = Western roan population A, WRB = Western roan population B, WRC = Western roan population C, SRA = Rest of Africa roan population A, rest of Africa population B, HYB = known hybrids, PTH = putative hybrids.

Genetic diversity for each population is summarized in Table 5. Overall, the genetic diversity in the Western roan populations is higher compared to populations from the East, Central and Southern African ESU, notwithstanding smaller sample sizes. The mean number of alleles (A) ranged from 4.15 - 6.07 and 4.26 - 5.70, while allelic richness (AR) ranged from 3.17 - 4.18 and 2.97 - 3.17 in the reference West African group, and East, Central and Southern African roan groups respectively. Observed heterozygosity (H_o_) in the Western roan group ranged from 0.67 - 0.72 and unbiased heterozygosity (H_z_) from 0.65 - 0.71 while H_o_ in the East, Central and Southern African roan varied from 0.57 - 0.63 and H_z_ from 0.605 - 0.609. The P_ID_ for the 27 loci was 5.5^-20^, thus the estimated probability of any two individuals by chance alone sharing the same mulitlocus genotype is 1.8^19^ for the 27 loci combined.

**Table 5.** Genetic diversity estimates for roan antelope (*Hippotragus equinus*).

| Samples | No. of samples | Mean no. of alleles per locus (A) | Allelic Richness (AR) | Unbiased Heterozygosity ($H_z$) | Observed Heterozygosity ($H_o$) | Inbreeding coefficient ($F_{IS}$) |
| --- | --- | --- | --- | --- | --- | --- |
| Western roan population A | 12 | 4.926 | 3.418 | 0.652 | 0.673 | -0.018 |
| Western roan population B | 14 | 6.074 | 4.182 | 0.714 | 0.709 | -0.022 |
| Western roan population C | 6 | 4.148 | 3.165 | 0.667 | 0.719 | -0.125 |
| Rest of Africa population A | 80 | 5.704 | 2.970 | 0.605 | 0.570 | 0.016 |
| Rest of Africa population B | 18 | 4.259 | 2.834 | 0.609 | 0.634 | -0.091 |
| Known hybrids | 8 | 4.963 | 3.425 | 0.671 | 0.598 | --- |
| Putative hybrids | 15 | 6.296 | 3.889 | 0.688 | 0.692 | --- |

## Discussion

An increasing number of species experience dramatic declining population numbers globally, with ample evidence suggesting that we are entering a mass extinction event. Although the drivers of these population declines are numerous and varied, the underlying root cause inevitably stems from anthropogenic pressures. Not surprisingly, hybridization and admixture of groups with distinct evolutionary trajectories are increasing, raising concerns about the integrity of a large number of species, especially those that experience disproportionately large human interest. For roan antelope, one of Africa’s most spectacular large antelope species, this is certainly the case. Although roan antelope numbers are increasing in South Africa (largely because of protection under private ownership), real concerns exist about their genetic integrity given admixture with West African roan antelope, also for export to neighbouring countries. We discuss our results here, and provide some suggestions for roan antelope conservation in South Africa.

### Evidence of hybridization

Using a suite of variable and informative microsatellite markers, we provide unequivocal evidence of hybridization and introgression between roan antelope naturally occurring in South Africa (East, Central and Southern African origin), and those of West African decent (a separate evolutionary significant unit; see [16]). More problematic, the identification of first and second generation backcrosses with *q*-values close to threshold values strongly suggest that hybrid individuals are viable and fertile; as also suggested from anecdotal evidence from some game farms. Although genetic diversity estimates were moderately higher in the known and putative hybrid individuals, it has previously been reported that F2 hybrids can display reduced fitness as a result of disruption of sets of co-adapted gene complexes by recombination [52,53], thereby weakening the entire gene pool of naturally occurring individuals. Our marker set was able to accurately identify F1 and F2 hybrids, as well as non-admixed individuals at thresholds of *q* = 0.20 and *q* = 0.15. However, the accurate classification of further backcrosses was less accurate at these thresholds (40% to 83%) with backcrossed individuals being incorrectly classified as non-admixed. The use of higher thresholds (*qi* = 0.10 and *qi* = 0.05) did increase the number of individuals correctly classified as backcrosses, however, this also resulted in an increase in the number of non-admixed individuals being incorrectly classified as hybrids. Thus in certain instances, backcrossed and double backcrossed individuals extend beyond the detection power of the current microsatellite marker panel.

The minimum number of markers required to accurately and consistently identify backcrosses is currently being debated. Simulation analysis in the grey wolf (*Canis lupus*) that hybridizes with domestic dogs (*C. lupus familiaris*) indicated that simply increasing the number of microsatellite markers used does not equate to an increase in the number of correctly identified admixed individuals [54]. It may be important to evaluate single nucleotide polymorphisms (SNPs) with high discriminating power to increase the ability to detect backcrossed and double backcrossed individuals, but in all likelihood thousands of SNPs may be required. Notwithstanding, the marker set described here represents the first step in assessing hybridization in roan antelope, and in the identification of hybrid individuals.

### Conservation management

As signatories to the Convention on Biological Diversity, South Africa has an obligation to conserve the genetic integrity of its biological diversity. Furthermore, admixture between distinct wildlife subspecies is prohibited under national and provincial legislation. Within South Africa, wildlife can be privately owned. There has been some debate about the legal rights of an owner to act in a certain manner with its property, and whether farming with wildlife should be managed and regulated any differently than, for example, agricultural stock such as cattle. Notwithstanding, current international, national and provincial legislation is clear in prohibiting admixture, irrespective of ownership.

The private ownership of biological diversity has been advantages for a large number of species, and the high commercial value attached to many of these species has undoubtedly aided in their conservation and protection; to the point where a number of species are doing better under private ownership compared with in protected areas or national parks [55]. Roan antelope is a prime example, but others include sable antelope, white and black rhinoceros, and bontebok to name but a few. Unfortunately, many of these species are intensively managed, with selection for specific desired traits. These management practises have unintended consequences, notably a loss of genetic diversity. In our study, a number of loci showed deviations from HWE and linkage disequilibrium; all which can be ascribed to small numbers of founding individuals and genetic drift on farms [56] which may, in the long term, compromise local adaptation [57]. To fully understand the impact that farming practises, notably intensive management and selection, have on wildlife populations, comparisons need to be done with naturally occurring populations on nature reserves.

Currently, the full extent of hybridization in South Africa between roan antelope belonging to the two distinct ESUs is unknown. Laboratory screening for permitting purposes (to either sell, or translocate animals) suggest that the occurrence of widespread introgression is low, and largely confined to specific game farms.

Animals of West African decent are no longer maladapted to South African conditions and have, over the span of 20 years, adapted to local conditions. The question that needs consideration is whether South Africa should safeguard the genetic integrity and genetic variability of both roan ESUs. If historic occurrence is considered, then all West African animals should be removed from South African populations. However, the South African situation has spawned several *ex situ* breeding programmes and agreements and/or animals that could be allowed to be backcrossed to obtain some form of purity, over four or five generations. This might improve genetic variation within the national population, but may not be desirable given that the impact of hybridization on the South African roan full genome is not known. Thus, we recommend the implementation and continuation of strict genetic monitoring for hybridization in roan antelope in South Africa. With the microsatellite marker set described here, and using a threshold of *qi* = 0.15, it is possible to detect F1 and F2 hybrids prior to translocation, thereby reducing and ultimately eliminating Western roan antelope alleles in the indigenous roan gene pools. In addition, management of roan in South Africa would benefit from a national meta-population conservation plan to inform translocations and reintroductions and to effectively monitor genetic diversity and further hybridization events.

## Acknowledgments

Anna M van Wyk was supported as a doctoral candidate under the Professional Development Programme of the National Research Foundation (NRF) as part of NZG grant no UID93062.

## Author contribution statement

AMvW, DLD, AK, JPG & BJvV wrote the main manuscript. AMvW, PSM & ASK conducted the laboratory analysis. AMvW, DLD & BJvV conducted genetic analysis of the data. All authors reviewed the manuscript.

## Additional information

### Conflict of interest

The authors declare that they have no conflict of interest.

### Compliance with ethical standards

Ethical approval was obtained from the Animal Research Ethics Committee, University of the Free State, South Africa (UFS-AED2017/0010) and the NZG Research Ethics and Scientific Committee (NZG/RES/P/17/18). Samples were stored in the NZG Biobank and access for research use of the samples was approved under a Section 20 permit from the Department of Agriculture, Forestry and Fisheries, South Africa (S20BB1917).

### Data availability

All species specific primers developed here will be submitted to GenBank following acceptance of this manuscript.

---

S1 Fig. a) Probability (-LnPr) of K = 1 − 6 averaged over 5 runs. b) Delta K values for real population structure K = 1 − 6.

